# Analysis of phylogenetic relationships and genome size evolution of the *Amaranthus* genus using GBS indicates the ancestors of an ancient crop

**DOI:** 10.1101/085472

**Authors:** Markus G. Stetter, Karl J Schmid

## Abstract

The genus *Amaranthus* consists of 50 to 70 species and harbors several cultivated and weedy species of great economic importance. A small number of suitable traits, phenotypic plasticity, gene flow and hybridization made it difficult to establish the taxonomy and phylogeny of the whole genus despite various studies using molecular markers. We inferred the phylogeny of the *Amaranthus* genus using genotyping by sequencing (GBS) of 94 genebank accessions representing 35 *Amaranthus* species and measured their genome sizes. SNPs were called by *de novo* and reference-based methods, for which we used the distant sugarbeet *Beta vulgaris* and the closely related *Amaranthus hypochondriacus* as references. SNP counts and proportions of missing data differed between methods, but the resulting phylogenetic trees were highly similar. A distance-based neighbor joing tree of individual accessions and a species tree calculated with the multispecies coalescent supported a previous taxonomic classification into three subgenera although the subgenus *A. Acnida* consists of two highly differentiated clades. The analysis of the Hybridus complex within the *A. Amaranthus* subgenus revealed insights on the history of cultivated grain amaranths. The complex includes the three cultivated grain amaranths and their wild relatives and was well separated from other species in the subgenus. Wild and cultivated amaranth accessions did not differentiate according to the species assignment but clustered by their geographic origin from South and Central America. Different geographically separated populations of *Amaranthus hybridus* appear to be the common ancestors of the three cultivated grain species and *A. quitensis* might be additionally be involved in the evolution of South American grain amaranth (*A. caudatus*). We also measured genome sizes of the species and observed little variation with the exception of two lineages that showed evidence for a recent polyploidization. With the exception of two lineages, genome sizes are quite similar and indicate that polyploidization did not play a major role in the history of the genus.

## 1 Introduction

The *Amaranthus* genus has a world-wide distribution and harbors between 50 and 70 species. The taxonomic differentiation of these species has proven difficult because only few traits are suitable for this purpose despite a high phenotypic diversity. In addition, there is a high level of phenotypic plasticity and a propensity to form interspecific hybrids and hybrid swarms (Brenner et al., 2013; Greizerstein and Poggio, 1994; Wassom and Tranel, 2005). Fertile hybrids can be obtained in crosses of distant species from different subgenera (Trucco et al., 2005). This disposition for natural hybridization further complicates the taxonomic differentiation of species.

Several species in the genus are of high economic importance and they include grain and vegetable crops as well as invasive weeds (Costea and DeMason, 2001; Sauer, 1967). The three species *A. cruentus, A. hypochondriacus* and *A. caudatus* were prehistorically cultivated in North, Central and South America for grain production. Together with their wild relatives *A. hybridus* and *A. quitensis* they form the Hybridus species complex and the latter two species have been suggested as ancestors of the three grain amaranth species, but the domestication history of amaranth is still under debate (Kietlinski et al., 2014; Sauer, 1967). *A. tricolor* is cultivated as leaf vegetable in Africa and Asia, in addition to *A. cruentus*, *A. dubius* and *A. hybridus*, which are also used as vegetable crops. Both seeds and leaves are high in micronutrients with a favorable amino acid composition (Rastogi and Shukla, 2013) and are therefore promoted as valuable crops for cultivation outside their native ranges. Appropriate cultivation conditions and protocols for efficient crosses allow to establish breeding programs to achieve this goal by breeding improved varieties of grain amaranths (Stetter et al., 2016). Weedy amaranths are the other group of economically and agronomically important species in the genus. The best known is Palmer amaranth (A. *palmeri*) because of its tolerance of the herbicide glyphosate. For example, yield losses in soybean fields due to Palmer amaranth infestation can range from 30 to 70 % (Bensch et al., 2003; Davis et al., 2015). Other weedy species of the genus include *A. tuberculatus, A. rudis* and *A. retroflexus*, which also lead to substantial yield losses in a diversity of crops (Bensch et al., 2003; Steckel and Sprague, 2004). The genus harbors several species that were reported to be resistant against herbicides (www.weedscience.com) and are useful models for studying the evolution of herbicide resistance.

The taxonomy and phylogeny of the genus was investigated using phenotypic traits and genetic markers. The most recent taxonomic revision defined three subgenera that include *Amaranthus Albersia, Amaranthus Acnida* and *Amaranthus Amaranthus* (Costea and DeMason, 2001; Mosyakin and Robertson, 1996). Previous studies with different genetic marker systems could not identify a consistent phylogeny of the genus that corresponds with the taxonomic classification (Lanoue et al., 1996; Chan and Sun, 1997; Wassom and Tranel, 2005; Das, 2014). Due to the difficulty of differentiating *Amaranthus* species by phenotypic traits, a total number 70 named species may be an overestimate if different populations of the same or closely related subspecies as well as hybrids are classified as different species. Almost 40 species are currently stored in the US (USDA/ARS) and German (IPK Gatersleben) *ex situ* genebanks and are readily available for taxonomic and phylogenetic analyses. A phylogeny of these species based on genome-wide genetic markers has the potential to improve the taxonomic classification and evolution of the whole genus beyond the grain amaranths and their close relatives, which are currently the best studied species (Jimenez et al., 2013; Xu and Sun, 2001). The rapid development of sequencing technology allows to utilize genome-wide polymorphisms from different species for phylogenetic analysis. Reduced representation sequencing methods, such as genotyping by sequencing (GBS) can provide thousands of single nucleotide polymorphisms (SNPs) for genetic analysis (Elshire et al., 2011; Poland et al., 2012) although for non-model species SNP detection can be challenging without a reference genome. In such species SNPs are identified by using the reference sequence of a different, but closely related species (Maughan et al., 2009), or the *de novo* assembly of sequencing reads (Catchen et al., 2011, 2013). Despite these limitations, GBS and related RADseq approaches have been used for phylogenetic analyses of both closely and distantly related taxa (Ariani et al., 2016; Eaton and Ree, 2013; Harvey et al., 2016; Nicotra et al., 2016)

Several software tools were developed for phylogenetic analyses based on biallelic markers. For example, SNAPP (SNP and AFLP Package for Phylogenetic analysis) infers species trees directly from biallelic markers by implementing a full multispecies coalescent model (Bryant et al., 2012). It integrates over all possible trees instead of sampling them explicitly, which results in a high statistical power, but is computationally expensive because it scales with the number of samples and markers (Paul et al., 2013).

The availability of a phylogenetic tree for a taxon allows to test hypotheses regarding phenotypic traits or other characters of interest. Species in the genus *Amaranthus* show variation in several traits such as reproductive system (monoecious vs. dioecious) and genome duplication. The latter process is commonly observed in plants and the genus *Amaranthus* is no exception because it is considered to be a paleoallotetraploid with a genome duplication between 18.4 and 34.0 Ma ago (Clouse et al., 2016). Haploid chromosome numbers reported for *Amaranthus* species are 16 and 17 (Greizerstein and Poggio, 1994, http://data.kew.org/cvalues), which indicates a cytological stability within the genus although there are several tetraploid species like *A. dubius* and *A. australis*, which likely have a different genome size or structure. Therefore, the variation of genome size within a genus is an interesting trait for analysis in the context of species formation and other phenotypic or ecological traits.

In this study we inferred the phylogeny of the genus *Amaranthus* using molecular markers and analyzed genome size variation to identify putative polyploidization events that may have played a role in speciation or influenced ecological traits. Of particular interest was the relationship of cultivated amaranths with their ancestors because the domestication history is not well understood. A genus-wide phylogeny may identify the ancestors of this ancient crop and allow to consider the evidence in the light of previous domestication models. Furthermore, knowledge of the evolutionary relationship between weedy amaranth species and their relatives allows to investigate the evolution of herbicide resistance. Previously, a diversity of molecular methods were used to infer a phylogeny of the *Amaranthus* genus that include seed proteins, RAPDs, AFLPs and SSRs (Chan and Sun, 1997; Khaing et al., 2013; Kietlinski et al., 2014). Most of these studies were applied to a subset of the species of the genus and gave inconsistent results (reviewed by Trucco and Tranel, 2011). In this study, we inferred a molecular phylogeny using a significantly larger number of species than previous studies using thousands of genome-wide markers identified with GBS. To evaluate the robustness of the phylogenetic analysis we compared different SNP calling methods that rely on reference sequences of distant relatives or on a *de novo* assembly of sequenced regions.

## 2 Material and Methods

### 2.1 Plant material

We obtained a total of 94 accessions representing 35 *Amaranthus* species from the USDA/ARS genebank and the German genebank at IPK Gatersleben (Table 1). Plants were grown under controlled conditions in standard gardening soil before leaves of young plantlets were collected for DNA and cell extraction. For genome size measurements all accessions were grown in two independent replicates.

**Table 1:** List of samples included in this study

| D | species | accession number | Genebank | Country |
| --- | --- | --- | --- | --- |
| 1 | *A. acanthochiton* | PI 632238 * | USDA/ARS | USA |
| 2 | *A. acutilobus* | PI 633579 | USDA/ARS |  |
| 3 | *A. albus* | PI 608029 | USDA/ARS | USA |
| 4 | *A. arenicola* | PI 667167 | USDA/ARS | Mexico |
| 5 | *A. asplundii* | PI 604196 * | USDA/ARS | Ecuador |
| 6 | *A. blitoides* | PI 649301 | USDA/ARS | USA |
| 7 | *A. blitum* | PI 490298 | USDA/ARS | Kenya |
| 8 | *A. blitum* | PI 612860 | USDA/ARS | USA |
| 9 | *A. californicus* | PI 595319 | USDA/ARS | USA |
| 15 | *A. caudatus* | PI 511680 * | USDA/ARS | Argentina |
| 26 | *A. caudatus* | PI 642741 | USDA/ARS | Bolivia |
| 28 | *A. caudatus* | PI 649230 † | USDA/ARS | Peru |
| 31 | *A. caudatus* | PI 649235 † | USDA/ARS | Peru |
| 34 | *A. caudatus* | PI 511679 * † | USDA/ARS | Argentina |
| 47 | *A. caudatus* | PI 649217 † | USDA/ARS | Peru |
| 50 | *A. caudatus* | PI 511681 * † | USDA/ARS | Bolivia |
| 51 | *A. caudatus* | PI 649228 * | USDA/ARS | Peru |
| 58 | *A. caudatus* | PI 608019 | USDA/ARS | Ecuador |
| 64 | *A. caudatus* | Ames 5302 † | USDA/ARS | Peru |
| 66 | *A. crassipes* | PI 649302 | USDA/ARS | USA |
| 67 | *A. crispus* | PI 633582 | USDA/ARS |  |
| 68 | *A. cruentus* | PI 511714 * | USDA/ARS | Peru |
| 76 | *A. cruentus* | PI 667160 | USDA/ARS | Guatemala |

| ID | Species | Accession number | Genebank | Country |
| --- | --- | --- | --- | --- |
| 80 | *A. cruentus* | PI 576481 | USDA/ARS | Mexico |
| 89 | *A. cruentus* | PI 433228 * † | USDA/ARS | Guatemala |
| 91 | *A. cruentus* | PI 658728 † | USDA/ARS | Mexico |
| 93 | *A. cruentus* | PI 511876 | USDA/ARS | Mexico |
| 101 | *A. cruentus* | PI 643037 † | USDA/ARS | Mexico |
| 103 | *A. deflexus* | PI 667169 | USDA/ARS | Argentina |
| 104 | *A. dubius* | Ames 25792 * | USDA/ARS | Panama |
| 105 | *A. fimbriatus* | PI 605738 | USDA/ARS | Mexico |
| 106 | *A. floridanus* | PI 553078 | USDA/ARS | USA |
| 107 | *A. graecizans* | PI 173837 | USDA/ARS | India |
| 110 | *A. hybr.* | PI 604571 † | USDA/ARS | Mexico |
| 119 | *A. hybr.* | PI 604564 † | USDA/ARS | Mexico |
| 120 | *A. hybr.* | PI 604566 † | USDA/ARS | Mexico |
| 123 | *A. hybridus* | Ames 5232 † | USDA/ARS | Peru |
| 127 | *A. hybridus* | PI 636180 | USDA/ARS | Colombia |
| 134 | *A. hybridus* | PI 667156 | USDA/ARS | Ecuador |
| 137 | *A. hybridus* | PI 604568 † | USDA/ARS | Mexico |
| 138 | *A. hybridus* | PI 604574 | USDA/ARS | Mexico |
| 140 | *A. hybridus* | Ames 5335 * | USDA/ARS | Bolivia |
| 141 | *A. hypochondriacus* | PI 649587 | USDA/ARS | Mexico |
| 146 | *A. hypochondriacus* | PI 633589 | USDA/ARS | Mexico |
| 158 | *A. hypochondriacus* | PI 604595 † | USDA/ARS | Mexico |
| 160 | *A. hypochondriacus* | PI 649529 | USDA/ARS | Mexico |
| 171 | *A. hypochondriacus* | PI 652432 | USDA/ARS | Brazil |
| 175 | *A. muricatus* | PI 633583 | USDA/ARS | Spain |
| 176 | *A. palmeri* | PI 633593 | USDA/ARS | Mexico |
| 177 | *A. polygonoides* | PI 658733 | USDA/ARS | USA |
| 178 | *A. quitensis* | PI 511747 | USDA/ARS | Ecuador |
| 185 | *A. quitensis* | PI 652426 | USDA/ARS | Brazil |
| 187 | *A. quitensis* | PI 652428 † | USDA/ARS | Brazil |
| 189 | *A. quitensis* | PI 652422 | USDA/ARS | Brazil |
| 192 | *A. quitensis* | PI 511736 * † | USDA/ARS | Bolivia |
| 196 | *A. quitensis* | Ames 5342 | USDA/ARS | Peru |
| 197 | *A. retroflexus* | PI 603852 | USDA/ARS | USA |
| 198 | *A. spinosus* | PI 500237 | USDA/ARS | Zambia |
| 199 | *A. standleyanus* | PI 605739 | USDA/ARS | Argentina |
| 200 | *A. tamaulipensis* | PI 642738 | USDA/ARS | Cuba |
| 201 | *A. tricolor* | PI 603896 | USDA/ARS |  |
| 202 | *A. tuberculatus* | PI 604247 | USDA/ARS | USA |
| 203 | *A. tuberculatus* | PI 603865 | USDA/ARS | USA |
| 204 | *A. tuberculatus* | PI 603872 | USDA/ARS | USA |
| 206 | *A. tuberculatus* | Ames 24593 | USDA/ARS | USA |
| 207 | *A. viridis* | PI 654388 | USDA/ARS | USA |
| 208 | *A. wrightii* | PI 632243 | USDA/ARS | USA |
| 209 | *A. spinosus* | AMA 13 | IPK |  |
| 210 | *A. crispus* | AMA 14 | IPK |  |
| 211 | *A. graecizans* | AMA 24 | IPK |  |
| 213 | *A. lividus* | AMA 49 | IPK |  |
| 216 | *A. graecizans* | AMA 62 | IPK |  |
| 217 | *A. acutilobus* | AMA 63 | IPK |  |
| 218 | *A. albus* | AMA 65 | IPK | Canada |
| 219 | *A. blitoides* | AMA 66 | IPK |  |
| 221 | *A. deflexus* | AMA 76 | IPK |  |
| 222 | *A. viridis* | AMA 79 | IPK | Peru |
| 223 | *A. dubius* | AMA 80 | IPK | Rwanda |
| 224 | *A. lividus* | AMA 87 | IPK | Rwanda |
| 225 | *A. powellii* | AMA 89 | IPK | Rwanda |
| 226 | *A. retroflexus* | AMA 93 | IPK | Mexico |
| 227 | *A. muricatus* | AMA 95 | IPK |  |
| 228 | *A. albus* | AMA 96 | IPK |  |
| 229 | *A. deflexus* | AMA 97 | IPK |  |
| 233 | *A. tricolor* | AMA 149 | IPK |  |
| 235 | *A. hybr.* | AMA 147 † | IPK |  |
| 238 | *A. retroflexus* | AMA 105 | IPK | China |
| 240 | *A. tricolor* | AMA 126 | IPK | Cuba |
| 242 | *A. dubius* | AMA 140 | IPK | Spain |
| 243 | *A. viridis* | AMA 175 | IPK |  |
| 244 | *A. powellii* | AMA 170 | IPK | Germany |
| 357 | *A. tucsonensis* | PI 664490 | IPK | USA |
| 360 | *A. australis* | PI 553076 | IPK | USA |
| 361 | *A. australis* | PI 553077 | IPK | USA |

\* Accessions not included in genome size measurements
† Accessions not included in SNAPP analysis

### 2.2 DNA extraction and sequencing

Genomic DNA was extracted with the Genomic Micro AX Blood Gravity kit (A&A Biotechnology, Poland) using CTAB extraction buffer for cell lysis (Saghai-Maroof et al., 1984). Double-digest genotyping by sequencing libraries (GBS) were constructed as described previously (Stetter et al., 2015). For each accession two samples with different barcodes were prepared to assure sufficient sequencing output per accession. Fragment sizes between 250 and 350 bp were selected with BluePippin (Sage Science, USA) and the resulting libraries were single-end sequenced to 100 bp on one lane of a Illumina HiSeq 2500 (Eurofins Genomics GmbH, Germany).

### 2.3 Data preparation and filtering

Raw sequence data were processed with a custom GBS analysis pipeline. First, reads were sorted into separate files according to their barcodes using Python scripts. Subsequently, read quality was assessed with fastQC (http://www.bioinformatics.babraham.ac.uk/projects/fastqc/). Due to lower read quality towards the end of reads, they were trimmed to 90 bp. Low quality reads were excluded if they contained at least one N (undefined base) or if the quality score after trimming was below 20 in more than 10% of the bases. Replicated data per accession were combined and subsequently analyzed as one sample.

### 2.4 *de novo* and reference-based SNP discovery

We used two different methods to call SNPs from the sequencing data, a *de novo* approach using Stacks 1.35 and an alignment to a reference genome. For the *de novo* approach we used the denovo_map.pl pipeline provided by Stacks to call SNPs directly from the processed data (Catchen et al., 2011, 2013). Highly repetitive GBS reads were removed in the ustacks program with option `-t`. Additionally, we analyzed data with two different minimum number of identical raw reads (`m = 3 and m = 7`) required to create a stack. These two settings resulted in different results in the SNP calling (Mastretta-Yanes et al., 2015) and we therefore used both settings for comparison. Two mismatches were allowed between loci when processing a single individual, and four mismatches between loci when building the catalog, which is the set of non redundant loci based on all accessions and is used as reference for SNP calling. SNPs were called with the Stacks tool populations 1.35 with filtering for different levels of missing values.

In addition to the *de novo* approach we used the sugar beet *(Beta vulgaris*) RefBeet-1.2 (Dohm et al., 2014) and the *Amaranthus hypochondriacus* draft genome (Clouse et al., 2016) as reference genomes to align sequence reads with bwa mem (Li and Durbin, 2009). SNPs were called with samtools 1.2 (Li et al., 2009). The resulting SNPs were filtered for different levels of missing values at a locus with vcftools (Danecek et al., 2011).

### 2.5 Phylogenetic analysis methods

We constructed a neighbor joining tree with 1000 bootstraps from the pairwise Euclidean distance between all 94 individuals based the four datasets using the R package ape (Paradis et al., 2004) and calculated an uncorrected neighbor joining network using the NeighborNet algorithm (Bryant and Moulton, 2004) with SplitsTree4 (Huson and Bryant, 2006).

We also used the multi-species coalescent implemented in SNAPP, which is part of the BEAST package, to infer species trees directly from unlinked biallelic markers (Bouckaert et al., 2014; Bryant et al., 2012). We reduced the number of individuals to a maximum of four per species because the SNAPP algorithm is computationally expensive. Additionally, we imputed the reference-map based datasets with beagle (Browning and Browning, 2016) before thinning all four datasets with vcftools (Danecek et al., 2011) to a distance of 100 bp which excludes multiple SNPs per GBS read. Since GBS loci are essentially randomly distributed throughout genome, we assumed that the assumption of unlinked biallelic markers was fulfilled after this filtering step. VCF files were converted to nexus format using a Python script and BEAST input files were created from these using BEAUti (Bouckaert et al., 2014). Mutation rates were calculated with BEAUti and default parameters were used for SNAPP. We conducted ten runs per dataset. Log files were analyzed with tracer 1.6 to examine convergence and converging log and tree files were combined using LogCombiner with 15% burn-in. The effective sample size (ESS) was adequate (> 200) for the important parameters but was lower for some *θ* values. We proceeded with the analysis as the low *θ* values should not influence the tree topology (Nicotra et al., 2016). TreeAnnotator was used to construct the ‘Maximum clade credibility’ tree and annotate it with posterior probabilities.

### 2.6 Genome size measurements and phylogenetic analysis

The genome sizes of 84 accessions representing 34 species were measured with flow cytometry and two independent replicates for each accession (Table 1). The tomato cultivar *Solanum lycopersicum* cv Stupicke was used as internal standard, due to its comparable genome size (DNA content = 1.96 pg; Dolezel et al., 1992). For the measurement, fresh leaves were cut up with a razor blade and cells were extracted with CyStain PI Absolute P (Partec, Muenster/Germany). Approximately 0.5 cm^2^ of the sampled leaf was extracted together with a similar area of the tomato leaf in 0.5 ml of extraction buffer. The DNA content was determined with CyFlow Space (Partec, Muenster/Germany) flow cytometer and analyzed with FlowMax software (Partec, Muenster/Germany). For each sample, 10,000 particles were measured. The DNA content was calculated as:

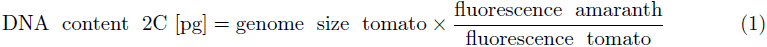

and the genome size (in Mbp) was calculated as:

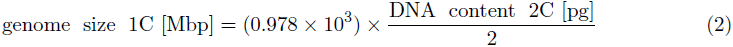

The conversion from pg to bp was calculated with 1 pg DNA = 0.978 × 10^9^ bp (Dolezel et al., 2003). Means were calculated using R software (R Core Team, 2014) and an ANOVA was performed to infer differences in genome size for the species.

We combined the genomic data with the genome size measurements to study the genome size evolution. The 1 C genome sizes (Mbp) were mapped on the phylogeny using parsimony reconstruction in Mesquite 3.04 (http://mesquiteproject.org). In addition we used the fastAnc function from the phytools R package to conduct a Maximum Likelihood reconstruction of ancestral states (genome sizes) with default parameters (Revell, 2012). For this analysis we inferred the genome size of *A. acanthochiton* as the mean between its two closest related species (*A. blitum* and *A. lividus*) because fastAnc does not allow missing values. A Brownian motion model implemented in the fastBM function in phytools (Revell, 2012) was used to simulate the random evolution of genome size over the tree. 1000 simulations were run and by using the distribution of genome sizes for each branch in the phylogeny the 0.25% and 97.5% were used to conduct a two-tailed test whether observed genome sizes were significantly smaller or larger than simulated sizes.

### 2.7 Data availability

Sequence reads were submitted to the European Nucleic Archive (ENA) under accession number PRJEB18745. Analysis scripts, aggregated sequencing data and genome size raw data are available under Dryad (http://datadryad.org/) DOI: http://dx.doi.org/10.5061/dryad.1bv83

## 3 Results

### 3.1 SNP discovery

Until reference genomes for any species can be created on a routine basis, methods like genotyping by sequencing (GBS) are an efficient method to survey genome-wide diversity in non-model species. To compare the use of GBS with and without a reference sequence for phylogenetic reconstruction of the *Amaranthus* genus, we used different methods and reference sequences for SNP calling. The number of aligned reads differed strongly between the *Beta vulgaris* and *Amaranthus hypochondriacus* references. Only 25.9% of the reads aligned to sugar beet and 74.8% to *A. hypochondriacus* (Table 2), which resulted in different SNP numbers. We identified 23,128 SNPs with the sugar beet and 264,176 SNPs with the *A. hypochondriacus* reference genomes. GBS data have a high proportion of missing values and the number of SNPs retained depends on the allowed proportion of missing values per SNP (Figure 1). For example, if no missing values are allowed only one SNP remained with the sugar beet and 247 SNPs with the *A. hypochondriacus* reference.

**Figure 1:**
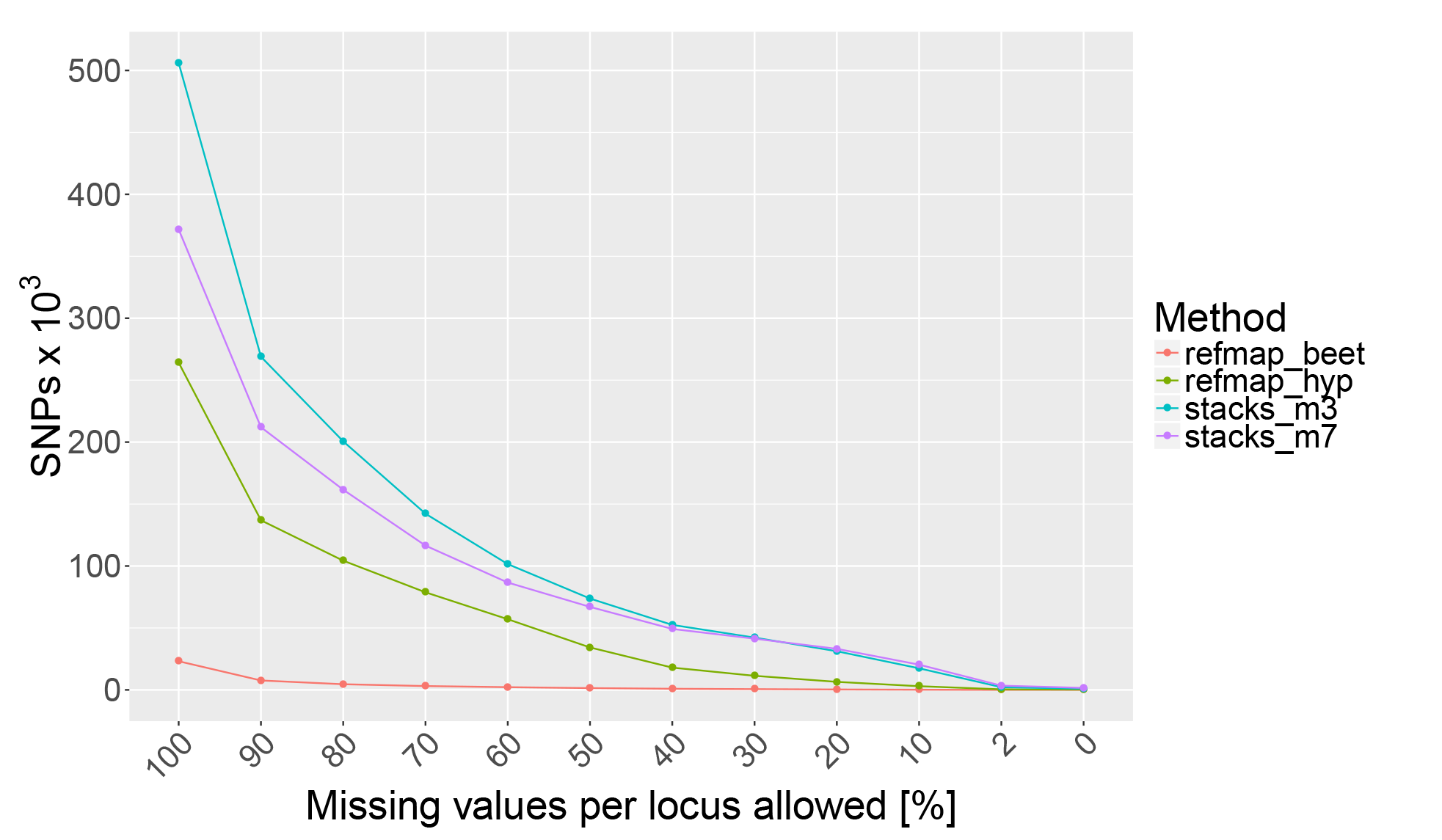
Number of SNPs recovered at di erent levels of missing values allowed per locus. Data sets are labeled as follows: refmap beet, reference mapping against sugar beet; refmap_hyp, reference mapping against *Amaranthus hypochondriacus*; stacks_m3, *de novo* assembly with Stacks using parameter value `m = 3` for minimal read coverage and stacks_m7, parameter value `m = 7`.

**Table 2:** Summary of four GBS datasets obtained by different SNP calling methods and parameters.

| Name | Reference map | Tool | Mapped reads | SNPs | Missing (%) |
| --- | --- | --- | --- | --- | --- |
| refmap_hyp | Ahypochondriacus_1.0 | BWA, Samtools | 166,935,845 (74.8%) | 2,978 | 5.2 |
| refmap_beet | RefBeet-1.2 | BWA, Samtools | 57,766,877 (25.9%) | 1,439 | 31.7 |
| stacks_m3 | *de novo* catalog | Stacks | 223,104,991 (100.0%) | 2,181 | 0.6 |
| stacks_m7 | *de novo* catalog | Stacks | 223,104,991 (100.0%) | 3,416 | 0.6 |

The *de novo* assembly with Stacks allowed us to use all reads for SNP detection at the cost that resulting contigs are unsorted and without position information on a reference genome. The minimum number of identical raw reads required to create a stack influences the SNP detection (Mastretta-Yanes et al., 2015). With a minimum number of three reads (m = 3) we obtained 505,981, and with seven reads (m = 7) 371,690 SNPs. After filtering out loci with missing values, m = 3 retained 949 and m = 7 retained 1,605 SNPs. The total number of SNPs recovered was higher for m = 3, but the number of SNPs without missing values was higher for m = 7. The two parameter values (m = 3 and m = 7) resulted in the same number of SNPs if a proportion of 20 to 30 % missing values per site were allowed. With both parameter values the *de novo* approach resulted in more SNPs before filtering than the reference-based SNP datasets (Figure 1). We were able to retain a large number of SNPs if missing data in one individual per GBS locus were allowed, which corresponds to a cutoff of 2% missing values (Figure 1). For the phylogenetic analysis of the reference-based datasets we allowed 10% (sugar beet reference) and 50% missing values (A. *hypochondriacus* reference). The resulting total number of missing values ranged from 0.6% for the *de novo* to 31.7% for the dataset based on the sugarbeet reference (Table 2). For the consecutive analyses we used all four datasets but in the following we present only the results obtained with the SNP data from the mapping against the *A. hypochondriacus* reference and include the other results as supplementary information because the results from all four data sets are very similar.

### 3.2 Phylogenetic inference

#### 3.2.1 Neighbor joining phylogeny

The neighbor joining phylogeny based on Euclidean distances of allelic states shows that most accessions cluster with other accessions from the same species (Figure 2). Within the Hybridus complex, however, there is no strong separation of the species into different clusters. Based on the species names, four clades are expected, but only three are observed. The first consists of *A. caudatus*, *A. quitensis* and *A. hybridus* that all originated from South America. The second clade includes *A. cruentus, A. hypochondriacus, A. hybridus*, which originated from Mexico, one *A. quitensis* accessions from Brazil and two hybrid accessions likely formed from species of the Hybridus complex. The third clade is formed by *A. cruentus, A. hypochondriacus* and *A. hybridus*, as well as two hybrids, and one *A. dubius* individual (242_dub; Figure 3). The accessions in this clade originated from Mexico, with the exception of two accessions of *A. cruentus* from Guatemala and one from Peru, and one *A. hypochondriacus* accession from Brazil. The NeighborNet network confirms this pattern and in addition outlines the extent of conflicting phylogenetic signals among accessions that may reflect gene flow or hybridization (Figure 3). The three accessions of the leaf vegetable amaranth *A. tricolor* cluster closely and form a clade with other *Amaranthus* species (Figure 2).

**Figure 2:**
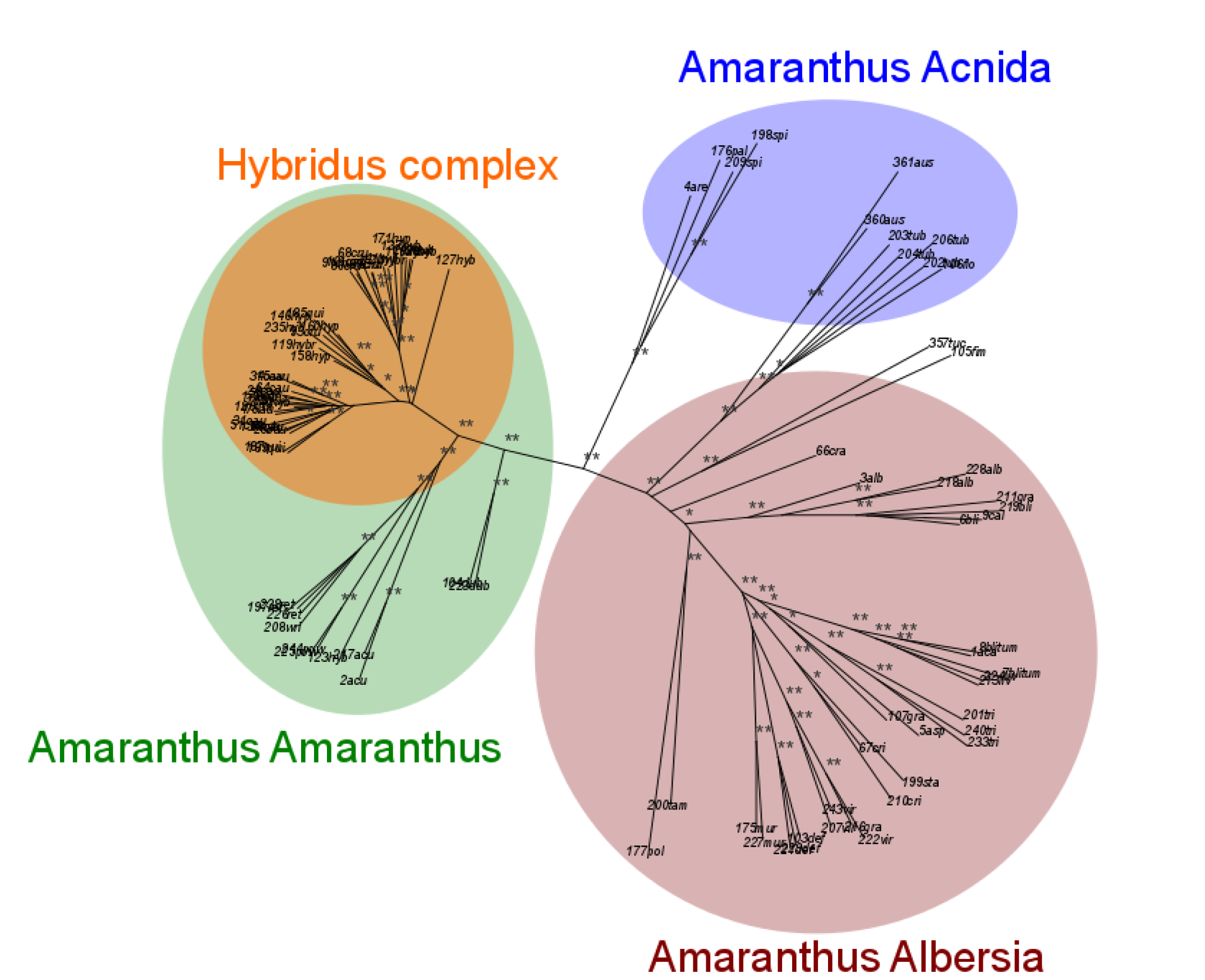
Neigbor joining tree calculated from the Euclidean distances of 94 individuals representing 35 *Amaranthus* species. Single stars (*) indicate bootstrap values over 90% and double stars (**) indicate bootstrap values of 100%.

**Figure 3:**
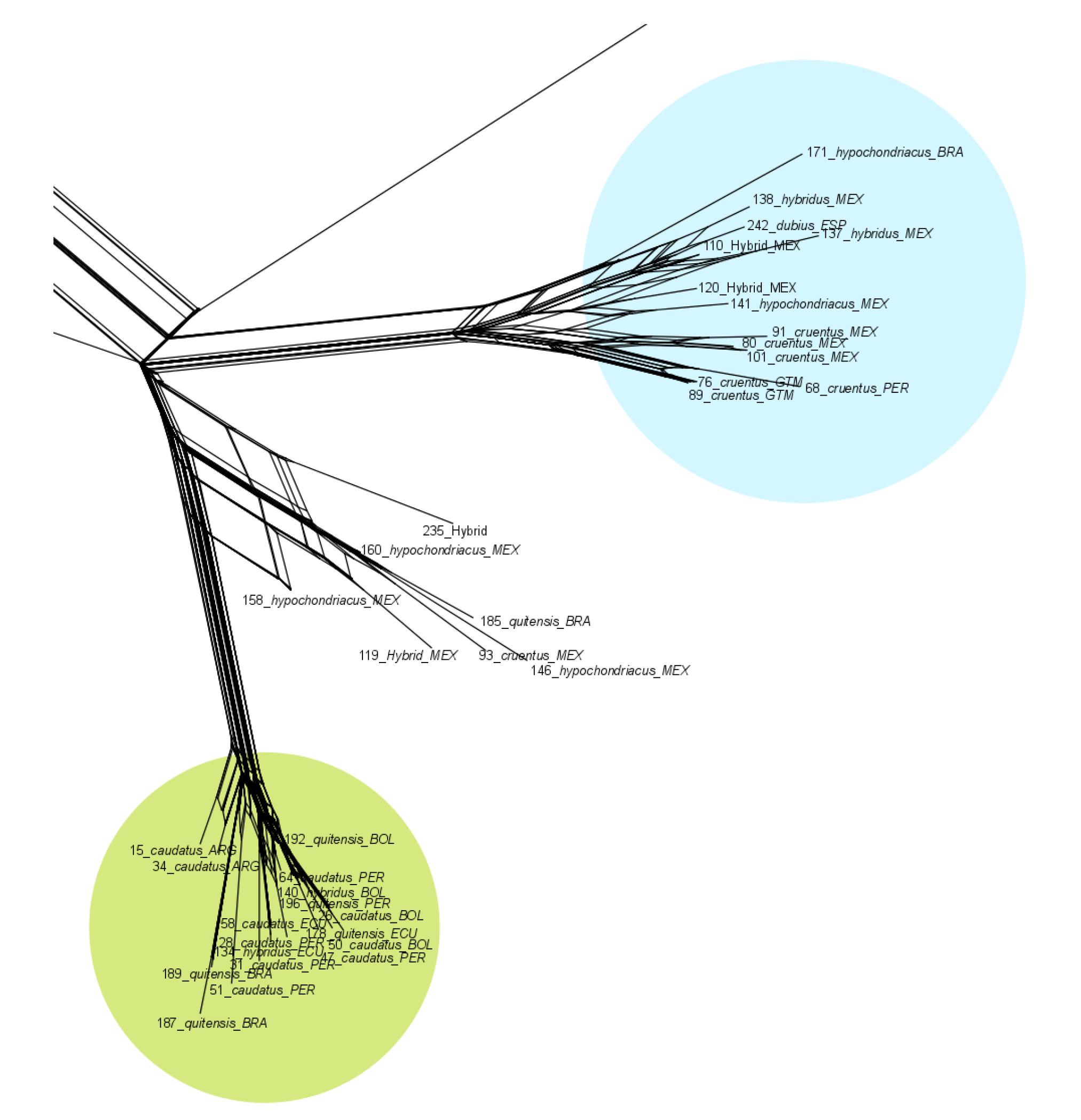
Section of the NeighborNet network showing the Hybridus complex. The blue circle includes the Central American grain amaranths (A. *hypochondriacus, A. cruentus*) and the close wild relative *A. hybridus*. The green circle includes South American grain amaranth (*A. caudatus* and the potential ancestors (A. *hybridus* and *A. quitensis*). The country of origin according to the genebank passport information is shown at the end of the name of each accession. The whole network is shown in supplementary figure S1.

Although the ability to resolve species level relationships seems to be limited with our data, the neighbor joining tree is consistent with the taxonomic classification into three subgenera that was previously defined using morphological traits (Figures 2 and S1). The phylogenies resulting from the four different SNP calling methods are highly similar and show that the tree topology of the genus is highly robust with respect to the SNP calling method (Figure S2).

#### 3.2.2 Phylogeny based on the multispecies coalescent

For inferring the phylogeny with the multispecies coalescent implemented in the SNAPP program we used a subset of individuals for two reasons. First, there were more individuals of the species from the Hybridus complex than of the other species which may bias the analysis, and second because the computation time scales exponentially with the number of individuals. Therefore we randomly sampled four individuals in those species with more than four genotyped accessions. The combined chain length without burn-in was 3,980,000 for the SNP data based on the *A. hypochondriacus* reference. The cloudogram derived from the SNAPP analysis allows to identify the degree of uncertainty for several clades in the tree (Figure 4). For the group of species that include *A. tricolor* and *A. crispus* there was a high uncertainty between the species. Within the Hybridus complex the uncertainty was high among the cultivated *A. caudatus* and its putative wild ancestors *A. quitensis* and *A. hybridus*. In contrast, the split between these three South American species and the Central American species *A. cruentus* and *A. hypochondriacus* was strongly supported. Overall, the Hybridus complex is well separated from the other species (Figure 4 and 5).

**Figure 4:**
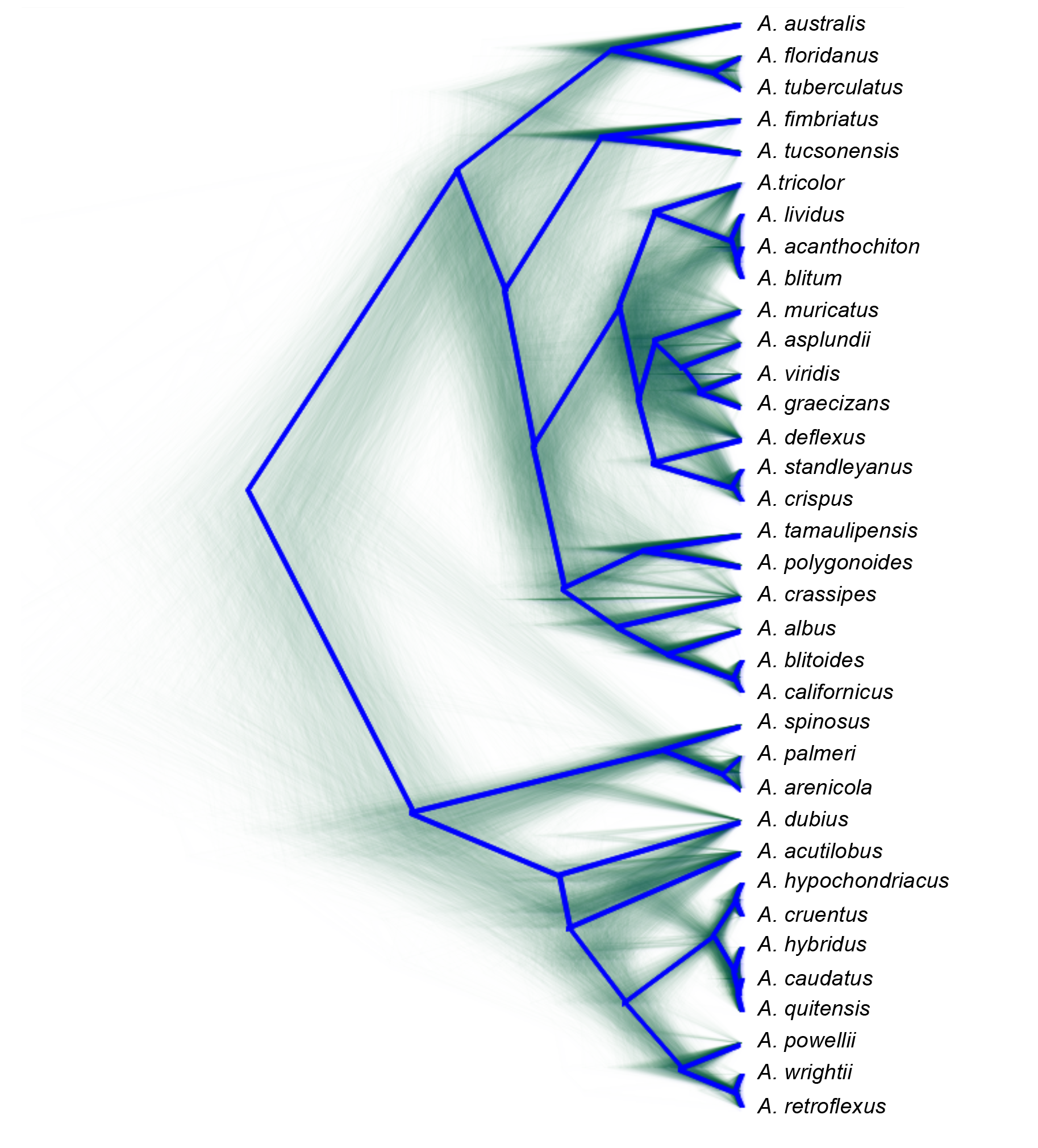
Species tree of *Amaranthus* based on the multispecies coalescent calculated with SNAPP. The cloudogram (green lines) represents 3980 individual trees and the consensus tree is shown in blue color.

### 3.3 Genome size evolution

The genome size measurements differed among the *Amaranthus* species although the range of variation was quite narrow (Table 3). Palmer amaranth has the smallest genome with a size of 421 Mbp, and *A. australis* the largest genome of 824 Mbp, which about twice the size of Palmer amaranth. Most species including the Hybridus complex had a genome size close to 500 Mbp (Table 3).

**Table 3:** Estimated genome size of *Amaranthus* species. n is the number of genotypes per species.

| species | $n$ | Size (Mbp) | Standard Error | Lower CI | Upper CI |
| --- | --- | --- | --- | --- | --- |
| *A. acutilobus* | 3 | 532.5 | 34.3 | 463.8 | 601.2 |
| *A. albus* | 3 | 530.3 | 33.4 | 463.2 | 597.3 |
| *A. arenicola* | 1 | 438.6 | 57.1 | 323.9 | 553.3 |
| *A. asplundii* | 1 | 535.0 | 57.1 | 420.2 | 649.7 |
| *A. australis* | 2 | 824.2 | 44.4 | 735.7 | 912.8 |
| *A. blitoides* | 3 | 521.9 | 33.4 | 454.8 | 588.9 |
| *A. blitum* | 2 | 748.8 | 40.6 | 667.2 | 830.4 |
| *A. californicus* | 1 | 547.9 | 57.1 | 433.2 | 662.6 |
| *A. caudatus* | 6 | 502.0 | 24.0 | 453.6 | 550.4 |
| *A. crassipes* | 1 | 512.5 | 62.4 | 388.1 | 637.0 |
| *A. crispus* | 2 | 576.0 | 40.6 | 494.4 | 657.6 |
| *A. cruentus* | 5 | 510.9 | 26.1 | 458.3 | 563.6 |
| *A. deflexus* | 3 | 640.2 | 33.4 | 573.1 | 707.2 |
| *A. dubius* | 2 | 711.9 | 40.6 | 630.3 | 793.5 |
| *A. fimbriatus* | 1 | 527.2 | 57.1 | 412.5 | 641.9 |
| *A. floridanus* | 1 | 658.2 | 57.1 | 543.5 | 772.9 |
| *A. graecizans* | 3 | 541.0 | 33.4 | 473.9 | 608.0 |
| *A. hybr.* | 3 | 508.0 | 33.4 | 440.9 | 575.0 |
| *A. hybridus* | 5 | 503.8 | 26.1 | 451.1 | 556.4 |
| *A. hybridus* x *A. hypochondriacus* | 1 | 523.8 | 57.1 | 409.1 | 638.5 |
| *A. hypochondriacus* | 5 | 506.4 | 26.1 | 453.7 | 559.0 |
| *A. lividus* | 2 | 685.8 | 40.6 | 604.2 | 767.4 |
| *A. muricatus* | 2 | 729.6 | 40.6 | 648.0 | 811.2 |
| *A. palmeri* | 1 | 421.8 | 57.1 | 307.1 | 536.5 |
| *A. polygonoides* | 1 | 512.3 | 57.1 | 397.6 | 627.0 |
| *A. powellii* | 2 | 512.3 | 40.6 | 430.7 | 593.9 |
| *A. quitensis* | 4 | 501.1 | 29.6 | 441.5 | 560.6 |
| *A. retroflexus* | 3 | 555.6 | 33.4 | 488.6 | 622.7 |
| *A. spinosus* | 2 | 471.6 | 40.6 | 390.0 | 553.2 |
| *A. standleyanus* | 1 | 502.9 | 57.1 | 388.2 | 617.6 |
| *A. tamaulipensis* | 1 | 524.9 | 57.1 | 410.2 | 639.6 |
| *A. tricolor* | 3 | 782.7 | 33.4 | 715.7 | 849.8 |
| *A. tuberculatus* | 4 | 675.6 | 27.0 | 621.4 | 729.8 |
| *A. tucsonensis* | 1 | 510.4 | 57.1 | 395.7 | 625.1 |
| *A. viridis* | 3 | 543.1 | 33.4 | 476.1 | 610.2 |
| *A. wrightii* | 1 | 534.3 | 57.1 | 419.6 | 649.0 |

To test whether changes in genome sizes in the phylogeny reflect random evolution or nonneutral processes, we mapped the genome sizes to the phylogenetic tree obtained with SNAPP (Figures 5 and S3). There was a tendency for decreasing genome sizes within the *Amaranthus* subgenus, and a high variation of genome sizes within the *Acnida* subgenus because it included both the individuals with the smallest and largest genome sizes. Figure 5 further shows that *A. dubius* has a larger genome than the other species of the *Amaranthus Amaranthus* subgenus. Even though there were significant differences in genome size between species, the ancestral branches have wide confidence intervals and significantly differ in recent splits but not in early ones (Figures S4 and S5). The ancestral genome size was inferred by fastAnc as 569 Mbp, but with a large confidence interval of 416 Mbp to 722 Mbp that includes almost all empirical genome size measurements of the extant species. Using a Brownian motion model we tested whether genome sizes differed in individual branches of the phylogeny given the complete tree. Several branches in the tree differ from such a random process. The lineage leading to *A. tricolor* and *A. australis* show significantly larger genome sizes that suggest that polyploidiziation likely influenced the genome sizes of these species. In contrast, the lineage leading to the weed *A. palmeri* has a significantly smaller genome size. The two clades of the *A. Acnida* subgenus consist of three species each. They are not only strongly separated according to the molecular phylogeny but also show different average genome sizes.

**Figure 5:**
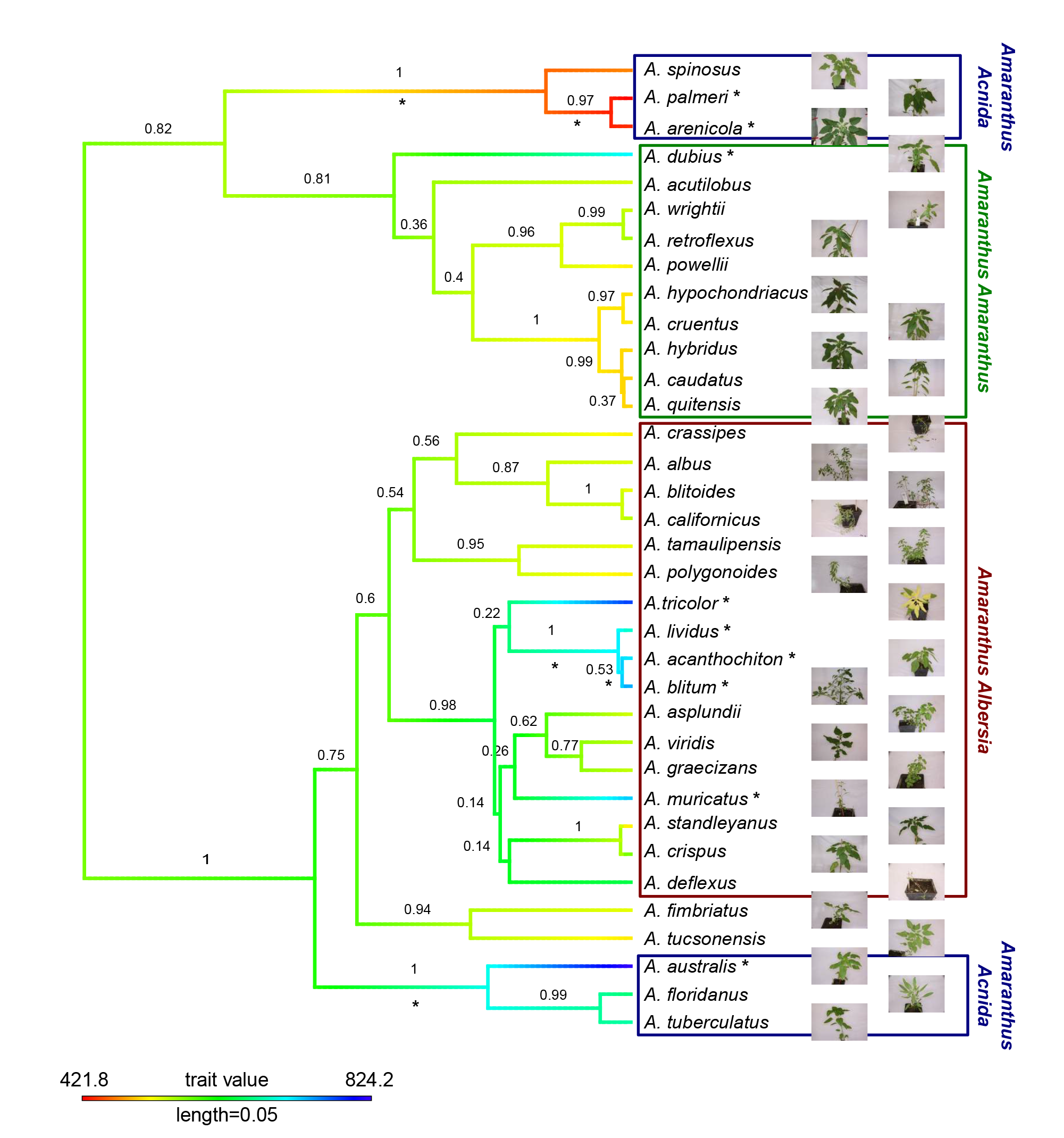
Genome size evolution mapped onto consensus tree obtained with SNAPP. The branch labels show posterior probabilities of genome size estimates of interior nodes obtained with a Maximum Likelihood method implemented in the fastAnc function of the phytools R package. Branch colors show estimated genome sizes in Mbp. Stars (*) indicate deviation from random evolution of genome size at 95% confidence level based on a two-tailed test. Group labels annotate taxonomic subgenera.

## 4 Discussion

### 4.1 Reference-based versus reference-free SNP calling

Genotyping by Sequencing (GBS) identifies thousands of markers but usually requires a reference sequence for mapping sequence reads. *De novo* methods allow to call SNPs without a reference genome. We compared both approaches to determine their efficiency in SNP identification. With the distant sugar beet genome as a reference only 26% of the sequencing reads could be used for SNP calling because the sequence divergence between sugar beet and *Amaranthus* species is too high for an efficient mapping despite the high synteny between *Amaranthus* and sugar beet (Clouse et al., 2016). This resulted in a small number of SNPs available for phylogenetic analysis. In contrast, the *de novo* assembly used all data and the number of SNPs obtained was even larger than from the mapping against the *A. hypochondriacus* genome. The proportion of missing data was also highest with the evolutionary distant sugar beet reference genome. Comparisons of different values for the number of identical reads (`-m` parameter) in Stacks showed that a smaller number of identical reads produced more SNPs, but we obtained more SNPs without missing values when requiring a larger number of identical reads, in accordance to earlier studies (Mastretta-Yanes et al., 2015). A reference genome from the same or a closely related species combines the advantage of a larger SNP number with linkage information (Andrews et al., 2016). Since the level of evolutionary divergence within the genus is unknown and only one reference sequence from an amaranth species was available, we compared the different approaches. Taken together, a comparison of the four SNP calling approaches with different numbers of SNPs and different levels of missing data showed that the resulting neighbor joining tree of the genus was quite robust with respect to SNP calling parameters, because it did not differ strongly between datasets (Figure S1). A major disadvantage of the *de novo* approach is that information about physical map positions of SNPs is missing and it can not be tested whether SNPs are unlinked. To increase the chance that SNPs are unlinked, which is a requirement of the SNAPP algorithm, we used a double-digest protocol for GBS and filtered for one SNP per GBS locus, which should allow the reconstruction of the phylogeny using the multispecies coalescent method (Andrews et al., 2016; Bryant et al., 2012; DaCosta and Sorenson, 2016). Such an approach was shown to be suitable for the phylogentic reconstruction of Australian *Pelargonium* using RADseq data (Nicotra et al., 2016).

### 4.2 Phylogeny of the whole *Amaranthus* genus

The species-rich genus *Amaranthus* has been divided into the three subgenera, *Amaranthus, Acnida* and *Albersia*. Several studies investigated species relationships in the genus using molecular markers, but most included only few species and did not allow conclusions for the whole genus (Chan and Sun, 1997; Lanoue et al., 1996; Kietlinski et al., 2014; Xu and Sun, 2001). We included all species that are currently available as *ex situ* conserved germplasm and genotyped several accessions per species to evaluate their evolutionary relationship (Figure 2). As expected, most accessions from the same species clustered together, and the subdivision of the genus into three subgenera based on phenotypic traits is largely consistent with our molecular data, although we observed some notable exceptions which we discuss below.

The species tree obtained with SNAPP largely reflects the neighbor joining tree which is based on individual accessions, but the cloudogram of all sampled species trees indicates uncertainties in the positioning of species like *A. deflexus, A. tricolor* and *A. crispus* in the tree topology (Figure 4). In contrast, a clustering of the genus into four basal clades is strongly supported (Figures 4 and 5). We compared our phylogeny with the published taxonomy of the *Amaranthus* genus (Mosyakin and Robertson, 1996). The subgenera *Amaranthus Amaranthus* and *A. Albersia* show a clear split at the root of the tree, but *A. Acnida* is split into two separate clades (Figure 5). The species of *A. Acnida* were categorized as dioecious and grouped based on this trait (Mosyakin and Robertson, 1996) although *A. palmeri* and *A. tuberculatus* were later described to be phylogenetically divergent (Wassom and Tranel, 2005). Another explanation for the observed split of *A. Acnida* species into two major groups may reflect the polyploid genomes of *A. tuberculatus, A. floridanus* and *A. australis* (see below). In our analysis, we treated all species as diploid and allowed only biallelic SNPs but polyploids may be characterized by high levels of heterozygosity (Figure S6) and harbor multiallelic SNPs, which are excluded from further analysis. Both factors may bias a phylogenetic inference. On the other hand, a high proportion of heterozygous loci would result in grouping the polyploid species in the same main branch as their ancestors or closest relatives. We conclude, however, that their grouping is correct because the posterior probabilities for the placement of these species in the phylogeny are very high.

### 4.3 Phylogenetic analysis of the Hybridus complex

The Hybridus complex contains the domesticated grain amaranths and putative ancestors such as *A. hybridus*. Previous studies suggested that the Hybridus complex comprises two clades (Adhikary and Pratt, 2015). We also identified the two clades, and in addition a third clade, which appears to be an intermediate of the other two other. It includes accessions from different species from Hybridus complex plus accessions that were labeled as ‘hybrids’ in the passport data and may have originated from interspecific hybridization. Interestingly, *A. hybridus* and *A. quitensis* accessions occur in all three clades (Figure 2), which may be explained by the geographic origin and geographic differentiation of these species. We previously suggested that *A. quitensis*, which is endemic to South America, and *A. hybridus* populations from the same region are a single species with a strong differentiation of geographically separated subpopulations within South America (Stetter et al., 2015). Since such a taxonomic grouping is still under debate and *A. quitensis* might be a separate subspecies of *A. hybridus*, we treated them as separate species in the phylogenetic analysis as was done before (Coons, 1978, 1982; Kietlinski et al., 2014). A comparison of the position of individual *A. hybridus* and *A. quitensis* accessions in the neighbor joining tree with the species tree obtained with SNAPP showed that in the former, the two species are not strongly differentiated from each other (Figure 2) whereas they form independent lineages in the species tree, but are closely related and in a monophyletic group with the three grain amaranths (Figure 5). This may be explained by the fact that SNAPP uses pre-defined groups which forces the algorithm to separate the species and therefore does not allow to evaluate whether *A. quitensis* can be considered as a separate species or is a subspecies with a high level of admixture.

The taxonomic interpretation of species relationships in the Hybridus complex is further complicated by the geographic origin of the accessions used in this study and by the effects of domestication. Sauer (1967) suggested that both *A. hybridus* and *A. quitensis* may have been involved in the domestication of the grain amaranths. Our analysis is consistent with this notion because the three grain amaranths *A. caudatus*, *A. cruentus* and *A. hypochondriacus* and their wild relatives *A. hybridus* and *A. quitensis* are separated from the other amaranths. The species tree suggests that both wild species are more closely related to the South American *A. caudatus* than to the Central American *A. cruentus* and *A. hypochondriacus*, but the neighbor joining tree of individual accessions splits *A. hybridus* accessions by their geographic origin and clusters *A. hybridus* accessions collected in South America with the South American *A. caudatus* and *A. hybridus* accessions collected in Central America with *A. cruentus* and *A. hypochondriacus*, which also are native to Central America (Figure 3).

Most evidence published so far suggests that *A. hybridus* is the direct ancestor of all three cultivated grain amaranth species (Chan and Sun, 1997; Kietlinski et al., 2014; Park et al., 2014; Stetter et al., 2015). *A. quitensis* is closely related to *A. caudatus* (Park et al., 2014; Xu and Sun, 2001; Stetter et al., 2015) and a low support of the split between *A. caudatus* and *A. quitensis* (Figures 4 and 5) reflects gene flow (Stetter et al., 2015) or indicates that *A. quitensis* is an intermediate between the wild *A. hybridus* and cultivated *A. caudatus* because it grows as weed in close proximity to grain amaranth fields and could have hybridized with *A. caudatus*. Another species for which a role in the domestication of grain amaranth was postulated is *A. powelli* (Sauer, 1967). In our analysis, as well as in a previous study, *A. powelli* is not closely related to the cultivated grain amaranths (Mallory et al., 2008) and therefore less likely a direct ancestor of *A. hypochondriacus* as proposed before (Park et al., 2014; Sauer, 1967; Xu and Sun, 2001).

Taken together, our analysis of the Hybridus complex is consistent with previous molecular phylogenies (Chan and Sun, 1997; Khaing et al., 2013) but we note that the GBS-based phylogenies show a weaker genetic differentation between the different species of the complex. In addition, both *A. caudatus* and *A. hypochondriacus* are more closely related to *A. hybridus* than to each other, which was observed before (Chan and Sun, 1997; Kietlinski et al., 2014). The *A. hybridus* accessions show a strong split along the North-South gradient (i.e., Central vs. South America), which supports the hypothesis that two different *A. hybridus* lineages were the ancestors of the three grain amaranths with a possible contribution of *A. quitensis* in the domestication of *A. caudatus* (Trucco and Tranel, 2011; Kietlinski et al., 2014; Adhikary and Pratt, 2015). Such a strong geographic pattern shows that in future studies requires a comprehensive geographic sampling to understand the evolutionary history of these species. Similar to the Hybridus complex other species of the genus should be sampled in greater detail to identify duplicated naming and issues with species delimitation. Species that are not yet included in germplasm collections should be collected and included in studies to determine the actual number of species in the genus.

### 4.4 Genome size evolution

The *Amaranthus* genes has undergone a whole genome duplication before speciation which was then followed by further duplication, chromosome loss and fusion events (Behera and Patnaik, 1982; Clouse et al., 2016). The mapping of genome size measurements onto the phylogeny revealed that the subgenus *Amaranthus* has a tendency towards smaller genomes, whereas species in the *Albersia* clade show increased genome sizes (Figure 5). These patterns are not strong and uniform within groups, however, because *A. dubius* has a larger genome size than expected for the clade. It may result from a genome duplication and a subsequent speciation of *A. dubius*, which is tetraploid (Behera and Patnaik, 1982). The genome size of *A. dubius* is not exactly twice the size of closely related species and indicates a loss of DNA after duplication. A similar pattern was observed in the genus Chenopodium which also belongs to the *Amaranthaceae* (Kolano et al., 2016).

Haploid chromosome numbers in the Hybridus complex species are variable. *A. cruentus* has 17, and the other species 16 chromosomes (Greizerstein and Poggio, 1994), although it does not seem to strongly influence genome sizes (Greizerstein and Poggio, 1994; Stetter et al., 2015, Table 3). For some species we observed a strong deviation in genome sizes from previously reported values. The genome sizes of *A. caudatus*, *A. cruentus* and *A. hypochondriacus* are within the previously reported range of 465 to 611 Mbp, but the genome sizes of *A. retroflexus, A. spinosus* and *A. tricolor* were about 200 Mb smaller than previous values. We also found that the five species of the Hybridus complex have similar genome sizes whereas previous measures from these species strongly differ from each other (Bennett and Smith, 1991; Bennett et al., 1998; Ohri et al., 1981, http://data.kew.org/cvalues). A strong variation in genome size was also observed in the dioecious *A. Acnida* subgenus. Previous molecular studies separated two members of this taxonomically defined subgenus *A. palmeri* and *A. tuberculatus* into different groups (Lanoue et al., 1996; Wassom and Tranel, 2005) and our phylogenetic analysis grouped the six species into two strongly separated clades of three species each, which differ by their average genome sizes. The genome size of *A. australis* is twice the size of *A. palmeri* and may result from a whole genome duplication (Mosyakin and Robertson, 1996). The closest relatives of *A. australis* are *A. florianus* and *A. tuberculatus*, which also have larger genome sizes than most species. This indicates that a polyploidization happened during the ancestral split of this group. In contrast, *A. palmeri* and its two closest relatives have the smallest genome sizes of the genus. The test for random evolution of genome size suggests that both clades deviate significantly from a model of random evolution due to independent instances of genome duplication and sequence loss (Figure 5).

Genome size may correlate with ecological and life history characteristics (Oyama et al., 2008). For example, one could postulate that herbicide tolerant weedy amaranths have a smaller genome size because they are faster cycling than their non-resistant relatives. We found that the genome sizes of the weedy amaranths in the different subgenera are highly variable and there does not seem to be a strong relationship between resistance and genome size. For other traits like mating system the number of species in the genus is currently too limited to allow strong conclusions regarding the evolution of the genome sizes. In addition to polyploidization, genome size evolution is also driven by transposable element (TE) dynamics (Bennetzen and Wang, 2014). Since GBS data sample only a small part of the genome and only one draft genome is currently available from the genus, it is not possible to evaluate the role of TEs in genome size evolution of the genus with these data.

## 5 Conclusions

GBS is a suitable approach for the phylogenetic analysis of the *Amaranthus* genus and allows a high taxonomic resolution. The large number of SNPs obtained from the *de novo* assembly of GBS sequencing reads and the high congruence of phylogenetic trees based on reference-mapping and *de novo* assembly indicates that a reference genome is not required and allows to study the molecular phylogeny of distantly related and non-model species. The inferred phylogeny based on 35 species largely confirms the previous taxonomic classification into three subgenera but also identified highly differentiated groups within the tree taxonomically defined subgenera. In particular, the subgenus *A. Acnida* consists of two strongly different groups with very different genome sizes, which may warrant a taxonomic revision. The comparison of a coalescent species tree with a distance-based tree of multiple individual accessions from each species identified clades in which gene flow, hybridization or geographic differentiation influenced the genomic relationship of species. The species in the Hybridus complex are closely related and were not separated along the species boundary, but are split into two main groups of accessions and species that reflect the geographically separated groups from South and Central America, respectively. The phylogeny of the genus further allowed to pinpoint the most likely ancestors and wild relatives of cultivated grain amaranths. In particular, *A. hybridus* appears to be the ancestor of all three crop amaranth species and the weed *A. quitensis* might be an intermediate between *A. hybridus* and *A. caudatus* or have contributed substantially to the domestication of *A. caudatus* by gene flow. The genome size measurements indicate that polyploidization events were rare in the genus. As in other plant taxa, further studies like the sequencing of the complete genomes of Amaranth species will be required to fully understand the relative importance of gene flow, hybridization and selection on the taxonomic relationships within the genus.

## Acknowledgements

We thank the USDA and IPK genebanks for providing the seeds used in this study and to David Brenner, Amaranth Curator of the USDA genebank for advice and discussion. We are grateful to Elisabeth Kokai-Kota for technical assistance in the laboratory. This work was supported by the F. W. Schnell Endowed Professorship of the Stifterverband für Deutsche Wissenschaft to K. J. S. We also acknowledge support by the state of Baden-Württemberg through bwHPC.

